# NanoTrans: an integrated computational framework for comprehensive transcriptome analysis with Nanopore direct RNA sequencing

**DOI:** 10.1101/2022.11.29.518309

**Authors:** Ludong Yang, Xinxin Zhang, Fan Wang, Li Zhang, Jing Li, Jia-Xing Yue

## Abstract

Nanopore direct RNA sequencing (DRS) provides the direct access to native RNA strands with full-length information, shedding light on rich qualitative and quantitative properties of gene expression profiles. Here with NanoTrans, we present an integrated computational framework that comprehensively covers all major DRS-based application scopes, including isoform clustering and quantification, poly(A) tail length estimation, RNA modification profiling, and fusion gene detection. In addition to its merit in providing such a streamlined one-stop solution, NanoTrans also shines in its workflow-orientated modular design, batch processing capability, all-in-one tabular and graphic report output, as well as automatic installation and configuration supports. Finally, by applying NanoTrans to real DRS datasets of yeast, *Arabidopsis*, as well as human embryonic kidney and cancer cell lines, we further demonstrated its utility, effectiveness, and efficacy across a wide range of DRS-based application settings.

## Introduction

The recent years have witnessed a rapid advancement of long-read sequencing technologies such as PacBio and Oxford Nanopore in terms of both read length and read accuracy. Combined with their single-molecular nature, this opens up exciting opportunities for many applications in both basic and applied biomedical research (Logsdon et al., 2020). Long-read-based RNA sequencing, best represented by Oxford Nanopore’s direct RNA sequencing (DRS), has also become increasingly popular, offering rich biological information while avoiding reverse transcription and amplification biases (Garalde et al., 2018). Accordingly, a number of dedicated bioinformatic tools have recently been developed for analyzing Nanopore DRS data with emphasis on specific applications, such as Mandalorion (Byrne et al., 2017) and Flair (Tang et al., 2020) for full-length isoform clustering, NanoCount (Gleeson et al., 2022) for expression quantification, nanopolish (Loman et al., 2015) and tailfindr (Krause et al., 2019) for poly(A) tail length estimation, EpiNano (Liu et al., 2019) and Xpore (Pratanwanich et al., 2021) for RNA modification, LongGF (Liu et al., 2020) and JAFFAL (Davidson et al., 2022) for gene fusion detection. While these tools greatly facilitated Nanopore DRS data analysis in their corresponding application scope, a unified framework that combines the power of these different tools are still badly needed. The Nextflow-based pipeline MasterOfPores is among the first to address this challenge (Cozzuto et al., 2020), but it lacks the support for important analysis such as differential gene/isoform expression detection, alternative splicing event evaluation, and fusion gene identification. More recent, a Snakemake-based pipeline named FASTdRNA (Chen et al., 2023) was developed with additional support for alternative splicing event evaluation, but still lacking important functionalities such as differential gene/isoform expression detection and fusion gene identification. Therefore, while these existing bioinformatic pipelines facilitated the wider adoption of Nanopore DRS, noticeable functional and practical limitations still exist in their current implementations.

In this study, we developed NanoTrans, an alternative solution for one-stop Nanopore DRS data analysis that cover diverse application scopes: expression quantification, isoform characterization, poly(A) tail length profiling, RNA modification evaluation, and fusion gene detection. As demonstrated with various real world testing examples and a head-to-head benchmarking comparison with MasterOfPores and FASTdRNA, NanoTrans shines in its rich functionality, robust stability, and computational optimization. Therefore, NanoTrans represents a step up in addressing the practical needs of the field, which helps to make the Nanopore DRS technology accessible to a broader research community for taking advantage of its full potentials.

## Results

### The framework design of NanoTrans

NanoTrans is a Linux-based computational framework for automated high-throughput Nanopore DRS data analysis. NanoTrans is self-contained, with all dependencies automatically installed and configured via a pre-shipped installer script. The design of NanoTrans is workflow-orientated, with a series of task-specific modules numbered according to their processing order (Figure 1 and Figure S1). Briefly, NanoTrans first performs Nanopore reads basecalling and reference genome preprocessing with its two starting modules numbered with “00”. The basecalled fastq reads are subsequently mapped to the preprocessed genome in a splicing-aware manner (module “01”), after which isoform clustering and quantification are further performed accordingly (module “02”). Based on the clustered and quantified isoforms, NanoTrans can perform different application-specific analysis such as isoform expression and splicing comparison (module “03”), isoform RNA modification identification (module “04”), isoform poly(A) tail length profiling (module “05”). In addition, reference-based gene fusion detection (module “06”) can be applied as well. A user-defined master sample table is used for specifying sample list, experimental design, and reads locations, based on which automatic batch processing and between group comparison are natively supported. Finally, an all-in-one final report containing all major tabular and graphic outputs will be generated for easy result exploration (module “07”).

**Figure 1.**
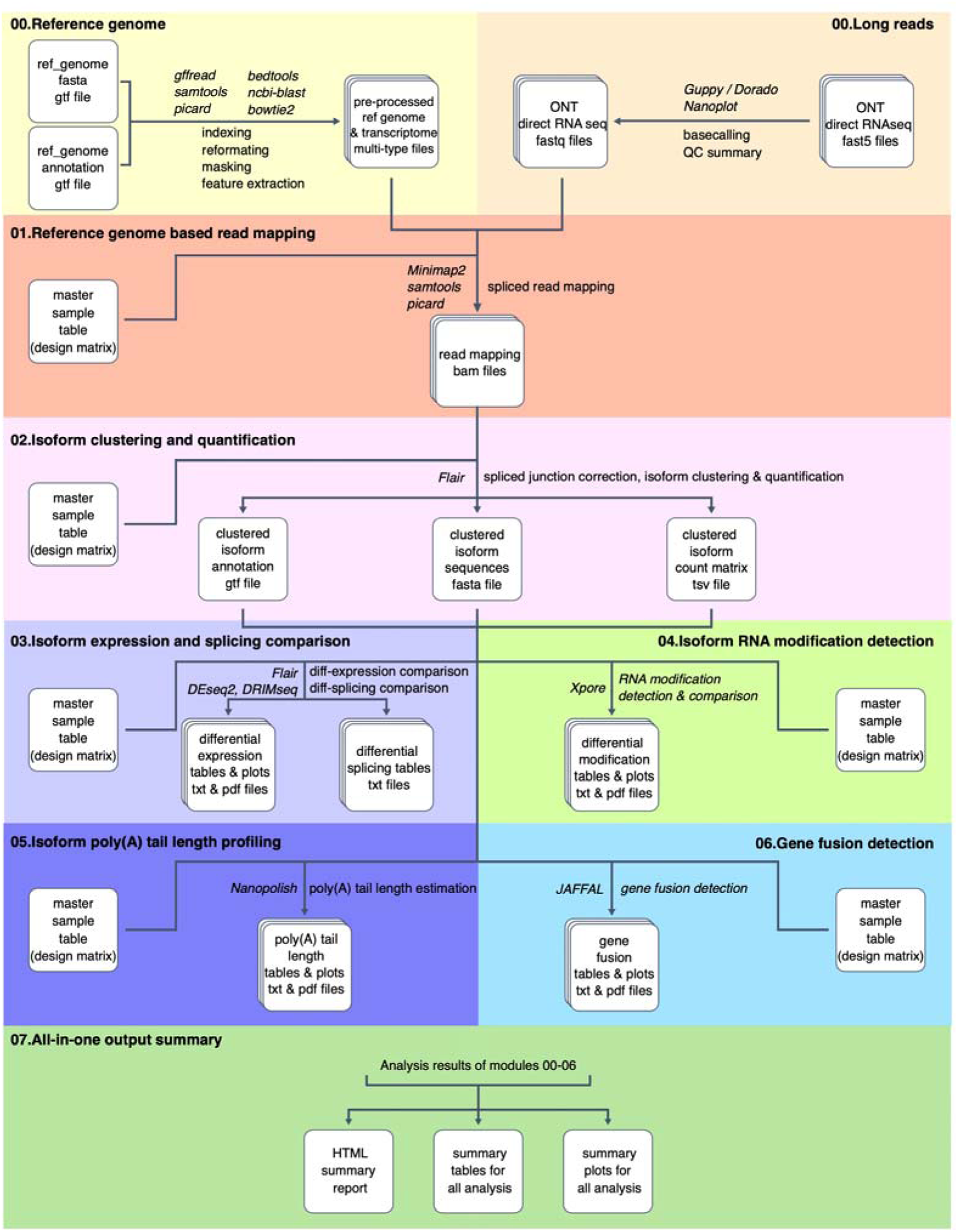
The framework design and application demonstration of NanoTrans. The overview of the NanoTrans framework. The names of third-party tools employed in each step are denoted in italics.

### Application demonstration summary of NanoTrans

To demonstrate the application of NanoTrans in real case scenarios, we first applied it to four public Nanopore DRS datasets (Table S1) from the budding yeast *Saccharomyces cerevisiae* (Dataset 1) (Tudek et al., 2021), the mustard plant *Arabidopsis thaliana* (Dataset 2) (Parker et al., 2020), the human embryonic kidney cell line (HEK293) (Dataset 3) (Chen et al., 2021), and the human lung adenocarcinoma (A549) and leukemia (K562) cancer cell lines (Dataset 4) (Gewartowska et al., 2021), respectively. These datasets were selected to demonstrate the performance of different functional modules of NanoTrans across different application settings (Table S2). With the yeast dataset (Dataset 1), we quantified the expression level and poly(A) tail length of each isoform and replicated the observation from the original study showing a negative correlation between RNA abundance and poly(A) tail length. With the *Arabidopsis* dataset (Dataset 2), we examined differential expression, differential splicing, and RNA modification. With the human embryonic kidney cell line dataset (Dataset 3), we verified the finding of the original study that TENT5A is responsible for the elongation of mRNA poly(A) tails. Finally, with the human cancer cell lines (Dataset 4), we identified a list of gene fusion alterations for the A549 and K562 cell lines respectively, highlighting the frequent chromosomal rearrangements in cancer genomes. Next, we present the performance of NanoTrans using the datasets above in details.

### Application demonstration of NanoTrans in *Saccharomyces cerevisiae*

Applying NanoTrans into the yeast dataset (Dataset 1) (Tudek et al., 2021), we quantified the expression level and poly(A) tail length of each isoform. We tried to test whether the outputs by NanoTrans could replicate the observation from the original study showing a negative correlation between RNA abundance and poly(A) tail length.

To do this, the *S. cerevisiae* reference genome (R64-1-1) and its associated feature annotation were retrieved from Ensembl (Table S1) and preprocessed with the 00.Reference_Genome module of NanoTrans for preparing various input files needed for downstream analysis. Then, the raw fast5 reads for the *S. cerevisiae* dataset were retrieved (Table S1) (Tudek et al., 2021) and provided to the 00.Long_Reads module of NanoTrans for basecalling and quality assessment. NanoTrans directly generates summarized tables and plots of this assessment, including read length, read quality, number of reads, read length N50 and so on (Table S3, Figure S2). Afterwards, the basecalled reads were mapped to the preprocessed reference genome via the 01.Reference_Genome_based_Read_Mapping module of NanoTrans. Given that this sequencing run contains admixed reads derived from both budding yeast and mouse samples, a stringent mapping quality filter of 20 (default: 0) was used at this step to filter out those non-yeast reads. Based on the read-mapping bam file, transcriptome clustering and quantification was further performed with the 02.Isoform_Clustering_and_Quantification module of NanoTrans. In the end, the clustered and quantified *S. cerevisiae* transcriptome isoform set was processed by the 05.Isoform_PolyA_Tail_Length_Profiling module for per-isoform poly(A) length estimation. By combining the quantification and poly(A) tail length profiling results, we examined their relationship. As reported by the original study that generated this Nanopore DRS dataset (Table S2) (Tudek et al., 2021), our analysis with NanoTrans recapitulated the negative relationship between the gene expression level and poly(A) tail length of each isoform (Figure 2A).

**Figure 2.**
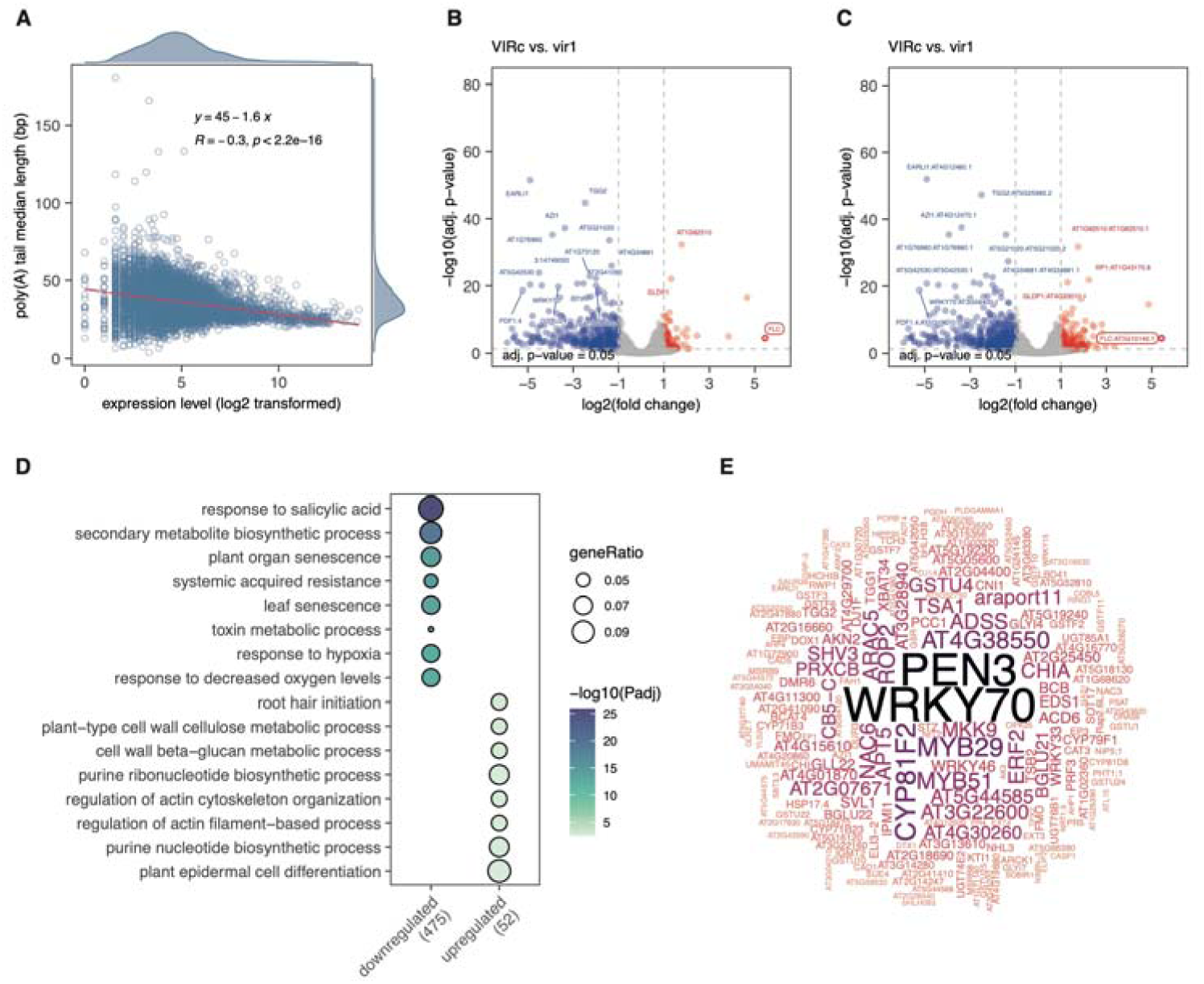
Applications of NanoTrans for isoform quantification and isoform expression comparison. (A) The negative correlation between isoform expression levels and poly(A) tail lengths identified in Dataset 1; the distribution of each isoform’s expression level (log2 transformed) is plotted along the x-axis while the distribution of each isoform’s median poly(A) length is plotted along the y-axis. (B-C) The differentially expressed genes (B) and isoforms (C) identified based on the comparison between the VIRc and *vir1* groups. False discovery rate (FDR) cutoff: 0.05. Fold change (log2 transformed) cutoffs: −1 and 1. (D) Biological processes enriched by differentially expressed genes (VIRc vs. *vir1*, adjusted p value < 0.05). (E) The size of words showed the frequencies of DEGs in the enriched GO terms; for example, *WRKY70* has the highest frequency among the enriched terms.

### Application demonstration of NanoTrans in *Arabidopsis thaliana*

To further demonstrating the performance of NanoTrans for analyzing differential expression, differential splicing, and RNA modification, we applied the corresponding modules in an *Arabidopsis thaliana* dataset (Dataset 2) (Parker et al., 2020). This dataset includes nanopore DRS reads of *VIRILIZER* loss of function mutant (*vir1*) and its full-function control (VIR complemented; VIRc), in which *VIRILIZER* is a conserved m^6^A writer complex component. Both the mutant and control lines had four biological replicates.

We first retrieved all the raw fast5 reads and used 00.Long_Reads module of NanoTrans for basecalling and quality assessment (Table S4). The basecalled reads were mapped to the preprocessed reference genome (TAIR10) via the 01.Reference_Genome_based_Read_Mapping module of NanoTrans with default settings. Based on the read-mapping bam file, transcriptome clustering and quantification was further performed with the 02.Isoform_Clustering_and_Quantification module of NanoTrans by leveraging all eight samples.

Next we used the 03.Isoform_Expression_and_Splicing_Comparison module of NanoTrans to perform differential expression and splicing analysis between the two comparison groups: *vir1* and VIRc. Since both comparison groups have ≥ 2 replicates, both gene-level and isoform-level differential expression can be assessed, and the batch effect correction is automatically taken care of behind scenes. The differentially expressed genes and isoforms are reported in tab-separated tables (sorted by p-value) and further visualized with volcano plots (Figure 2B-C). In such volcano plots, differentially expressed genes are labeled with “gene_name” only (Figure 2B), while differentially expressed isoforms are labeled with “gene_name:isoform_id” tags (Figure 2C). NanoTrans recaptured the differential gene expression signal on the *FLC* (*AT5G10140*) gene reported by the original study (Table S2), highlighting the important impact of the m^6^A machinery on flowering time regulation in *Arabidopsis*. More interestingly, we made a previously unreported finding that the *WRKY70* gene is among the strongly differentially expressed genes with the lowest p-values, suggesting its change in expression heavily linked to the interruption of the m^6^A machinery. Gene ontology (GO) enrichment of the differentially expressed genes further highlighted *WRKY70*’s prevalence across significantly enriched biological processes (Figure 2D-E). This finding is consistent with a recent report that m^6^A regulates the normal development in *Arabidopsis* by destabilizing the senescence-related transcripts, including *WRKY70* (Sheikh et al., 2024).

Regarding the differential splicing analysis, four types of alternative splicing events are examined with NanoTrans: intron retention (ir), alternative 3’ splicing (alt3), alternative 5’ splicing (alt5), and cassette exons (es). A summary table with differentially spliced isoforms (sorted by p-values) is reported. The detailed isoform splicing forms and usage across all samples can be further visualized using the NanoTrans.02.Plot_Isoform_Usage.sh script in the 02.Isoform_Clustering_and_Quantification module of NanoTrans. In Figure 3, we showed an example of such visualization, in which a novel isoform and two alternative splicing events were identified for the *Arabidopsis POM1* (*AT1G05850*) gene (Figure 3A). Samples from the two compared groups show considerable differences in their isoform usages, with the *vir1* group showing uniformly less usage for this novel transcript (Figure 3B). Furthermore, the complexity and alternative splicing of transcripts were also achievable through Module 02 of NanoTrans. For example, NanoTrans accurately identified the mutually exclusive exons between exon 2 and exon 3 in the *FLM* (*AT1G77080*) gene (Figure 3C). Secondly, it can precisely reveal highly complex transcript, such as AT1G48090.6 (Figure 3D). These two instances were also highlighted in the original study that produced this DRS dataset (Parker et al., 2020).

**Figure 3.**
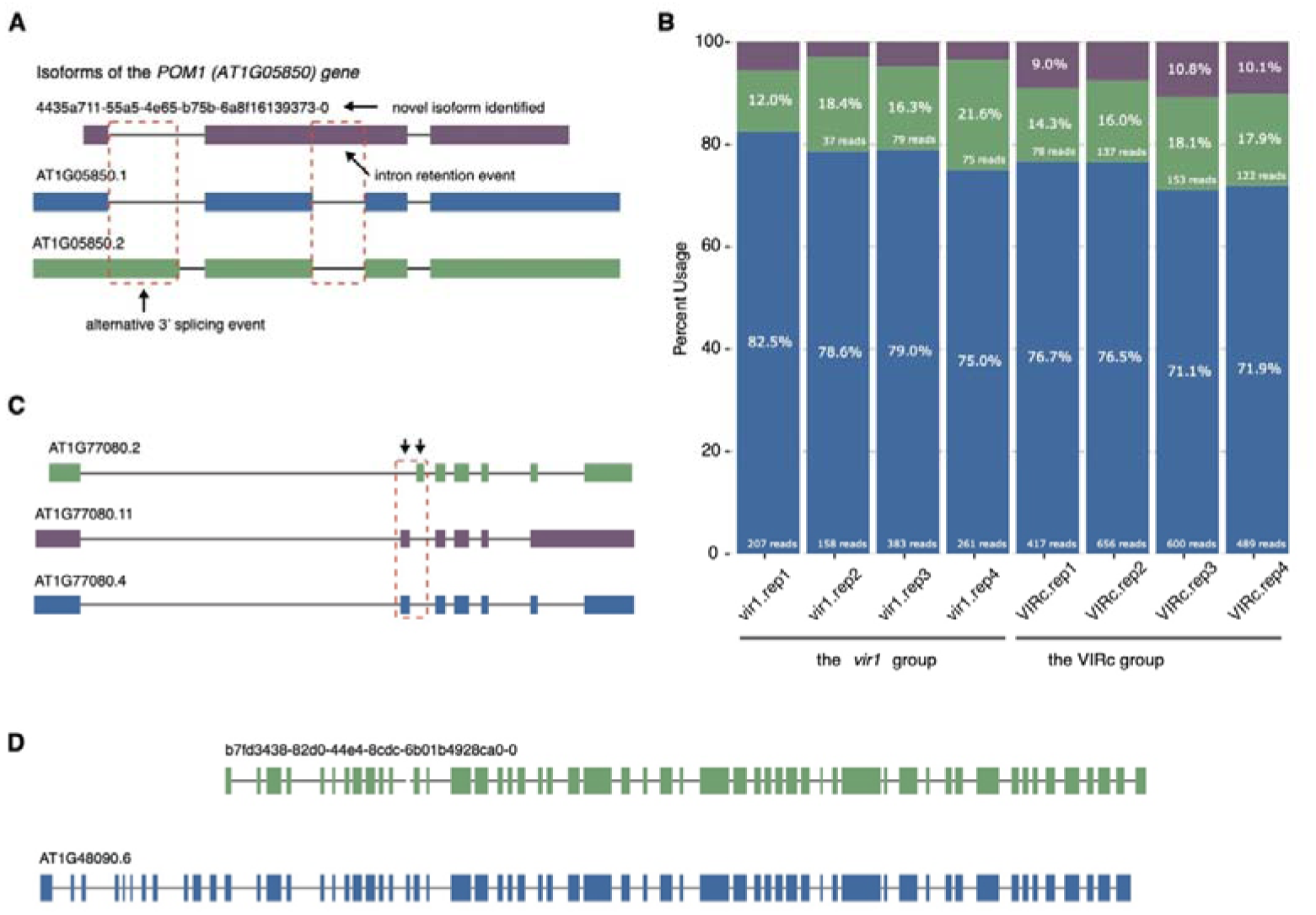
Examples of differentially spliced genes identified based on the comparison between the VIRc and *vir1* groups. (A) A novel transcript that is previously unannotated is identified for the *POM1* (*AT1G05850*) gene. The two identified intron retention events of this novel transcript are further highlighted based on its comparison to the two already annotated transcripts. (B) The different isoform usages of the three transcripts identified in panel A across all samples from the *vir1* and VIRc groups. (C) The mutually exclusive exons (exon 2 and exon 3, indicated by black arrows) in *FLM* (*AT1G77080*) gene is highlighted. (D) A complex transcript composed of a multitude of exons, AT1G48090.6, is identified.

We also used the 04.Isoform_RNA_Modification_Identification module of NanoTrans to evaluate transcript-specific kmers (5-mers) with differential RNA modification rates between the two defined comparison groups: *vir1* and VIRc (Figure 4A-B). Furthermore, the characteristics of differentially modified kmers can be explored and visualized using the script NanoTrans.04.Plot_characteristics_kmers.sh within the analysis module 04 of the NanoTrans framework. Specifically, the differentially modified kmers exhibited characteristics comparable to the original study, such as predominant distribution in 3’ untranslated regions (UTRs), a pronounced peak downstream of stop codons, and substantial overlaps between the identified kmers and polyadenylation sites (PASs) (Figure 4C-G). Taken together, these observations recapitulated the characteristics of differentially modified kmers identified in original study of this dataset (Parker et al., 2020).

**Figure 4.**
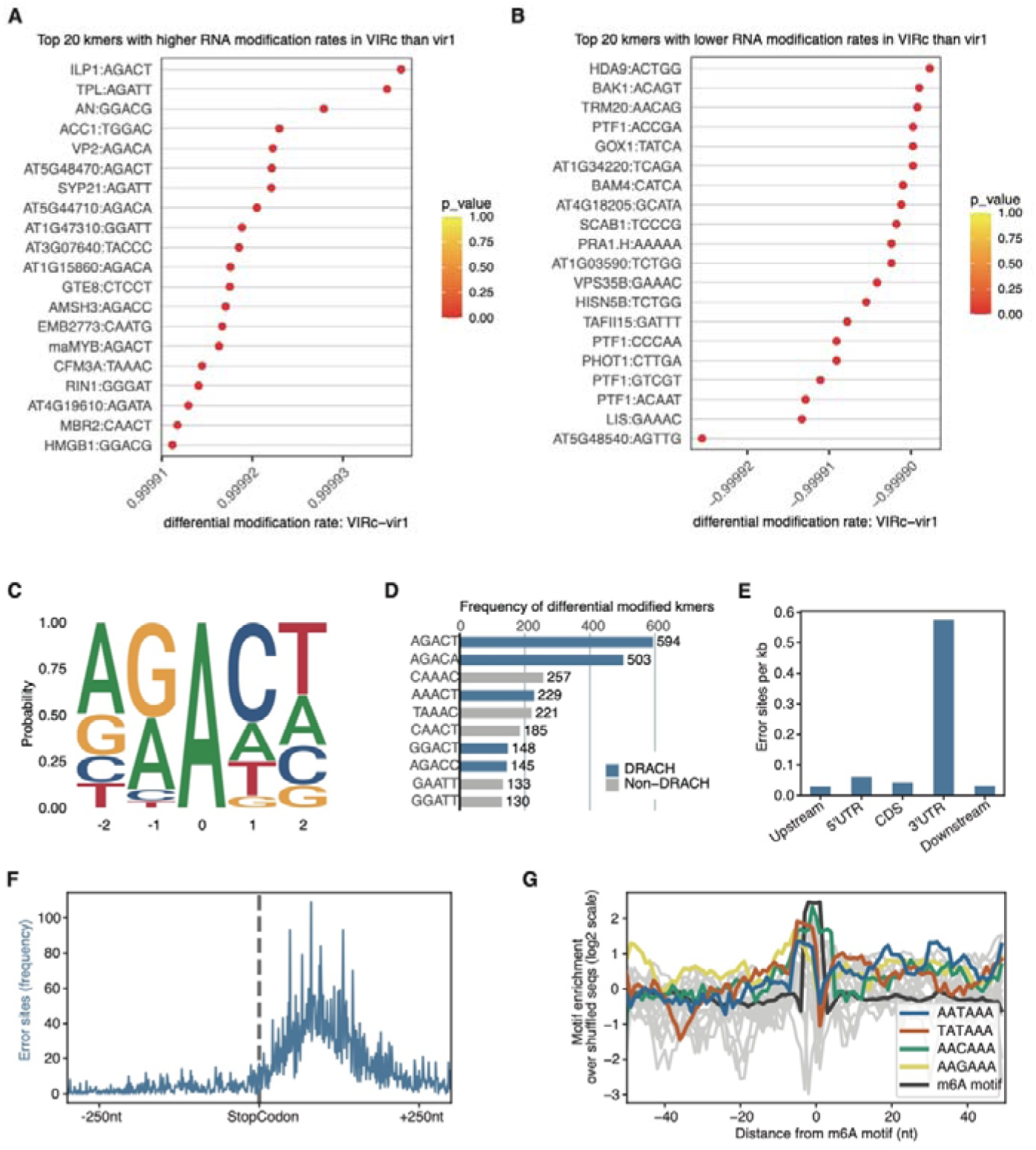
Characteristics of differentially modified kmers between the VIRc and *vir1* groups. The top 20 kmer-gene combinations with higher (A) or lower (B) RNA modification rates in the VIRc group than the *vir1* group. (C) The sequence logo showed the preference of filtered differentially modified kmers (DMKs, differential modification rates > 0.5, p-value < 0.05, and only kmers with center A were retained). (D) The top 10 DMKs with the highest frequency. (E) Distribution of the DMKs on distinct genomic features. (F) Frequency of DMKs around stop codons. (G) Overlaps between RNA modifications and poly(A) signal sites. NanoTrans directly outputs this kind of plot for each analyzed sample.

### Application demonstration of NanoTrans in human embryonic kidney cell line

Dataset 3 (BioProject accession: PRJEB39819; Reads accession: ERR5534379-ERR5534382) contains reads derived from two comparison groups (TENT5A^WT^ vs. TENT5A^MUT^) of the cell samples with the human embryonic kidney cell line HEK293 background (Gewartowska et al., 2021). The samples from the TENT5A^MUT^ group express a mutant version of the TENT5A protein with mis-sense mutations in its putative catalytic residues (D139N; D141N). There are two replicates in each comparison group. We named the two samples from the TENT5A^WT^ group as TENT5A.WT.rep1 and TENT5A.WT.rep2, while naming the two samples from the TENT5A^MUT^ group as TENT5A.MUT.rep1 and TENT5A.MUT.rep2. The decompressed raw reads were provided to the 00.Long_Reads module of NanoTrans for basecalling and quality assessment. The read length and quality summary of the basecalling results are shown in Table S5. The human reference genome (GRCh38) and its associated feature annotation were retrieved from Ensembl (release 107) and preprocessed with the 00.Reference_Genome module of NanoTrans. The basecalled reads were mapped to the preprocessed reference genome via the 01.Reference_Genome_based_Read_Mapping module of NanoTrans with default settings. Based on the read-mapping bam file, transcriptome clustering and quantification was further performed with the 02.Isoform_Clustering_and_Quantification module of NanoTrans by leveraging all four samples. The clustered and quantified human transcriptome isoform set was processed by the 05.Isoform_PolyA_Tail_Length_Profiling module of NanoTrans for isoform poly(A) tail length estimation.

First, we compared the poly(A) tail lengths of all isoforms between the two comparison groups: TENT5A^WT^ and TENT5A^MUT^. As Figure 5A and 5B show, the isoforms from the TENT5A^WT^ group showed slightly longer poly(A) tails in comparison to those from the TENT5A^MUT^ group across the whole genome (mean difference = 6.512; p-value =1.923×10^-82^). In addition to such genome-wide comparison, we also generated a selected gene list for genes that are associated with extracellular matrix constituents (GO: 0031012) by using the AmiGO2 web utility (http://amigo.geneontology.org). By further limiting our isoforms of interests to these genes, we saw much more pronounced differences in terms of poly(A) tail length between the two groups, with the genes from the TENT5A^WT^ groups showing longer poly(A) tails in general (mean difference = 11.705; p-value =1.646×10^-7^; Figure 5C-D). Such observation is consistent with the original study (Table S2) (Gewartowska et al., 2021).

**Figure 5.**
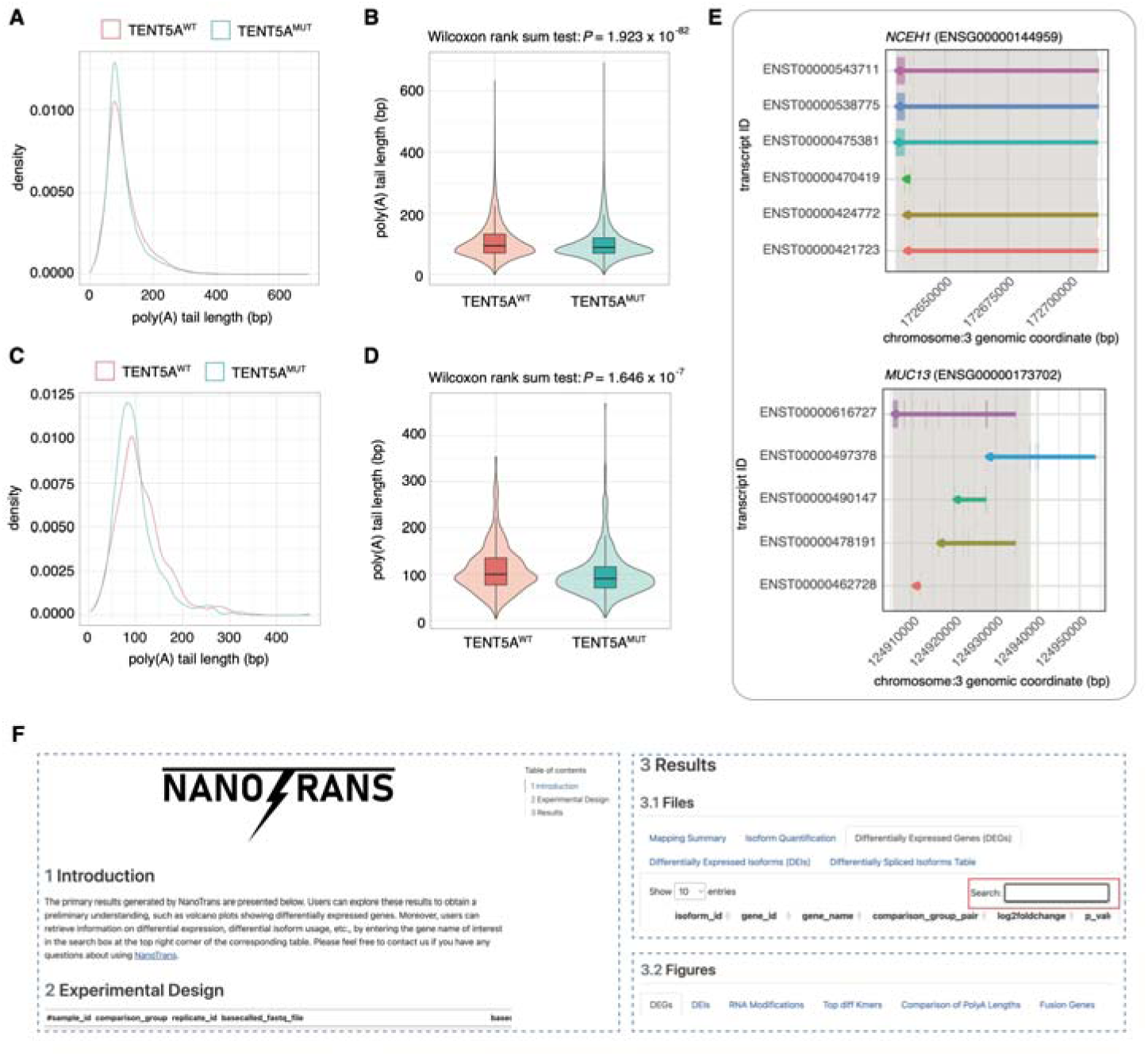
Applications of NanoTrans for poly(A) tail length estimation and gene fusion detection and its all-in-one summary report feature. (A) The density plot of genome-wide per-isoform poly(A) tail length distributions for the TENT5A^WT^ and TENT5A^MUT^ groups. (B) The violin plot of genome-wide per-isoform poly(A) tail length comparison between the TENT5A^WT^ and TENT5A^MUT^ groups. (C) The density plot of the per-isoform poly(A) tail length distribution of genes associated with extracellular matrix constituents (GO: 0031012) for the TENT5A^WT^ and TENT5A^MUT^ groups. (D) The violin plot of the per-isoform poly(A) tail length comparison of genes associated with extracellular matrix constituents (GO: 0031012) between the TENT5A^WT^ and TENT5A^MUT^ groups. (E) The *NCEH1*:*MUC13* gene fusion identified in the A549 human lung cancer cell line. The genome arrangements of all transcripts derived from the left donor (*NCEH1*) and right donor (*MUC13*) are presented with their corresponding transcription directions indicated by arrows. The genomic regions that constitute the fused transcript are highlighted in light grey. (F) An all-in-one HTML report that summarizing the results. The report contains three sections, in which the Results section gathered the primary files and figures generated by NanoTrans.

### Application demonstration of NanoTrans in human cancer cell lines

Dataset 4 (BioProject accession: PRJEB44348; Reads accession: ERR9286473 and ERR9286521) contains reads derived from two human cancer cell lines: the lung adenocarcinoma cell line A549 and the leukemia cell line K562 (Chen et al., 2021). We named these two samples as SGNex.A549.rep1 and SGNex.K562.rep1 respectively. The decompressed raw reads were provided to the 00.Long_Reads module of NanoTrans for basecalling and quality assessment. The read length and quality summary of the basecalling results are shown in Table S6. The human reference genome (GRCh38) and its associated feature annotation were retrieved from Ensembl (release 107) and preprocessed with the 00.Reference_Genome module of NanoTrans. We used the 06.Fusion_Gene_Detection module of NanoTrans to detect fusion genes based on the pre-processed reference genome and transcriptome by NanoTrans. For each sample, a summary table (sorted by the confidence level) was generated to report potential fusion genes and the associated transcript-based evidence.

Based on this dataset, we identified *NCEH1*:*MUC13* as a high-confidence fusion event for the A549 cell line while finding *BAG6:SLC44A4* and *GSE1:ATXN1-AS1* as high-confidence fusion event for the K562 cell line. These findings echoed well with the CCLE fusion gene call set (Ghandi et al., 2019)(Table S2). For each identified potential gene fusion event, NanoTrans also plots the genome arrangement of all transcripts associated with the left and right donor partners and highlights the corresponding genomic regions corresponding to the fused transcript. As an example, the *NCEH1*:*MUC13* gene fusion is graphically depicted, with the donor regions that make up the fused transcript highlighted in light grey (Figure 5E). This fusion should be triggered by an intrachromosomal rearrangement on chromosome 3.

### An all-in-one HTML report for result summary

Finally, the Module 07 provides an HTML report rendered by Quarto (https://quarto.org/), integrating essential files and figures to offer users a convenient exploration of the primary results. The report consists of three main sections: Introduction, Experimental Design, and Results (Figure 5F). Within the Results section, files and figures from each module are organized across different page tabs, providing a streamlined presentation. Moreover, users can easily locate a specific gene by entering its names into the search box at the top-right corner of the corresponding table.

### Benchmarking comparison between NanoTrans and similar existing tools

There are at least two bioinformatics tools, MasterOfPores (Cozzuto et al., 2020) and FASTdRNA (Chen et al., 2023), have been developed for Nanopore DRS data analysis. Hence, we compared these tools in terms of their functionality, ease of use, and computational efficiency.

Leveraged by functionality, NanoTrans, MasterOfPores, and FASTdRNA all covered the following functions in their latest implementations: 1) preprocessing (basecalling, quality control, reads mapping, reads counting); 2) RNA modification detection; 3) poly(A) tail length estimation (Table 1). In addition, NanoTrans and FASTdRNA have dedicate modules for alternative splicing detection in comparison to MasterOfPores. Most noticeably, only NanoTrans provides supports for differential gene/isoform expression detection and fusion gene identification analysis, which are important applications of DRS data (Table 1). In addition, while all three tools provide human-readable tabular results, NanoTrans stresses more on intuitive results summarization and visualization, which can greatly assist data interpretation towards better biological insights. Regarding ease of use, in addition to Linux command lines, MasterOfPores and FASTdRNA further require users’ familiarity with workflow system tools such as Nextflow and Snakemake. Additionally, the setup of MasterOfPores requires the pre-installation of containers such as Docker or Apptainer (formerly Singularity). These prerequisites raised extra challenges for end users. In comparison, NanoTrans only relies on Linux command lines, with a design philosophy to minimize the requirement of users’ bioinformatic knowledge when executing it. To gauge the computational efficiencies of NanoTrans, MasterOfPores, and FASTdRNA, we conducted a benchmarking comparison based on the same testing dataset (Dataset 5, reads quality shown in Table S7) for their commonly supported analysis, which include Nanopore reads preprocessing, RNA modification detection, and poly(A) tail length estimation. In comparison to NanoTrans, we found MasterOfPores took 18 additional hours to run through all analysis and consumed twice of much of disk space (Table 1). As for FASTdRNA, maybe because of its ongoing version updates, we cannot successfully run through its full workflow. With minor bug fixing, we were able to execute its preprocessing module (dRNAmain) and found both NanoTrans and MasterOfPores significantly outperform FASTdRNA in terms of computational efficiency (Table 1).

**Table 1.**
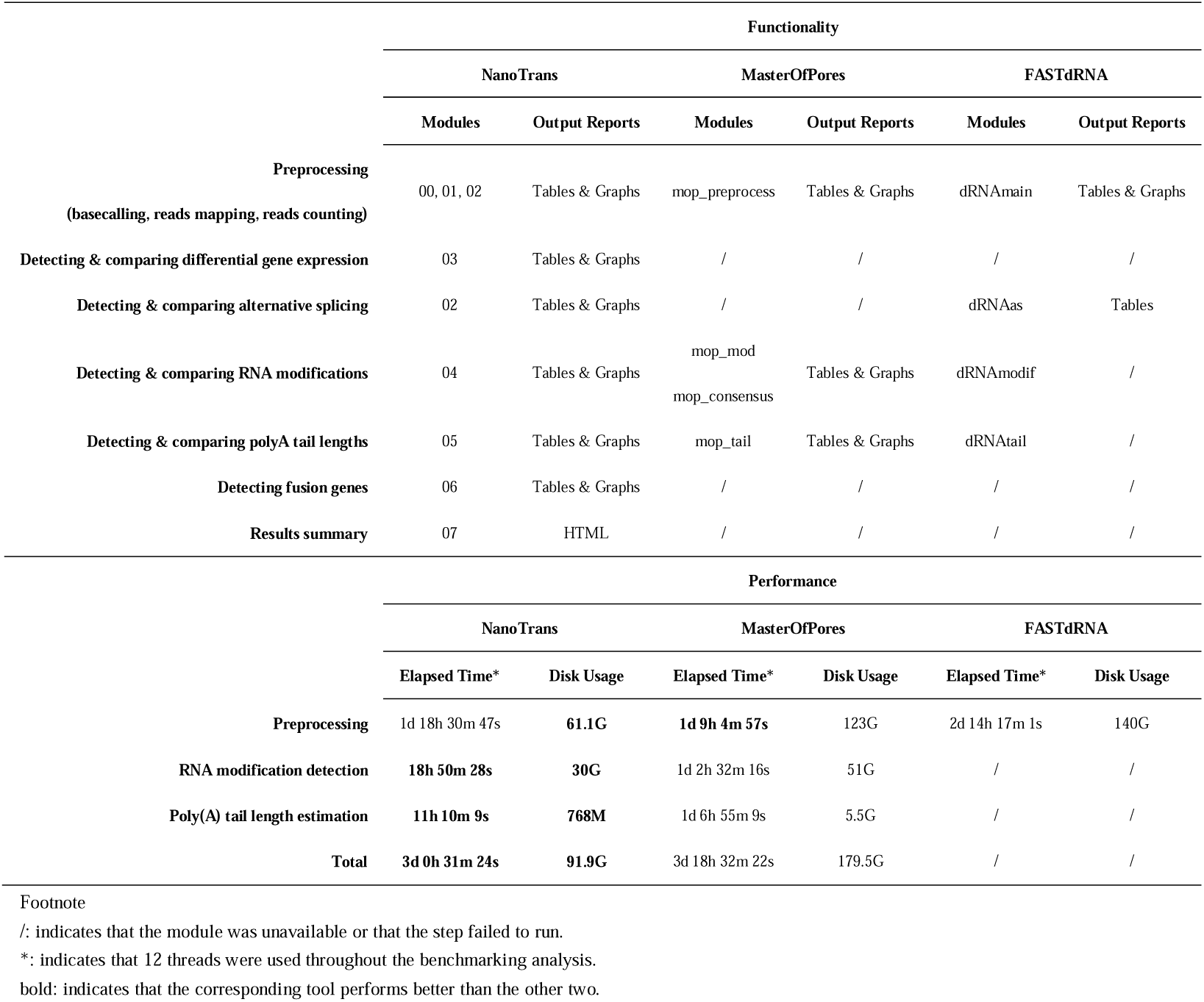
Comparison of NanoTrans, MasterOfPores, and FASTdRNA.

## Discussion

In this work, we established a framework named NanoTrans to facilitate easy and standardized Nanopore direct-RNA-sequencing (DRS) data analysis for not only experienced bioinformaticians but wet-lab experimentalists with beginners’ training on bioinformatics as well. NanoTrans alleviates the time and energy-consuming challenges of tools selection, parameter optimizing, workflow scripting, results visualization for DRS data analysis, offering a unified and smooth experience that enables the end users to better focus on biological insights. A detailed step-by-step manual with real world testing examples is further provided to guide the users through the workflow even when they have limited bioinformatics knowledge. Under the hood, NanoTrans comprises seven modules that comprehensively cover different aspects of DRS data analysis, ranging from the pre-processing of raw data, such as basecalling and mapping, to dedicated downstream analysis, such as identifying RNA modifications and gene fusion events. In most cases, executing NanoTrans modules only requires executing a single command to run. Finally, NanoTrans provides an all-in-one HTML report that allows users to explore results conveniently and comprehensively.

In this study, we demonstrated the robustness and versatility of NanoTrans by applying it to four diverse DRS datasets generated based on yeast, *Arabidopsis*, and human cell lines. Important conclusions of original studies that generated these DRS datasets were validated during our re-analysis with NanoTrans (Table S2). In addition, novel biological insights such as the importance of *WRKY70* gene expression change in response to the m^6^A machinery disruption is further revealed via NanoTrans during this process, which highlights the value of NanoTrans in facilitating biology-driven Nanopore DRS data analysis. Additionally, NanoTrans is designed and implemented to be as much user-friendly as possible, requiring minimal prior knowledge on Linux command lines from the end users. Technical complications of DRS data processing are automatically handled behind the scenes. Admittedly, Nanopore direct RNA sequencing and the best practice of its data analysis are both advancing very rapidly. In future, we will continue to maintain and update NanoTrans to keep it synchronized with the development of the field. The current implementation of the NanoTrans framework makes it very straightforward to incorporate newly developed tools when needed. We expected NanoTrans to grow with the field, continuously enabling researchers to make insightful biological discoveries via Nanopore DRS technology.

## Materials and methods

### Software prerequisites

NanoTrans is designed for a desktop or computing server running an x86-64-bit Linux operating system, preferably also with the multithreaded processor support. The installation process requires a stable internet connection and the following Linux software compilation prerequisites.

- bash (https://www.gnu.org/software/bash/)
- bzip2 and libbz2-dev (http://www.bzip.org/)
- gcc and g++ (https://gcc.gnu.org/)
- git (https://git-scm.com/)
- GNU make (https://www.gnu.org/software/make/)
- gzip (https://www.gnu.org/software/gzip/)
- libopenssl-devel (https://github.com/openssl/openssl)
- libcurl-devel (https://curl.se/libcurl/)
- java runtime environment (JRE) v1.8.0 (https://www.java.com)
- perl v5.12 or newer (https://www.perl.org/)
- tar (https://www.gnu.org/software/tar/)
- unzip (http://infozip.sourceforge.net/UnZip.html)
- wget (https://www.gnu.org/software/wget/)
- zlib and zlib-devel (https://zlib.net/)
- xz and xz-devel (https://tukaani.org/xz/)
- perl-devel (https://pkgs.org/download/perl-devel)
- R 3.6 or newer (https://www.r-project.org/)

### Software installation and configuration

A bash script is pre-shipped with NanoTrans to perform automatic installation and configuration for NanoTrans’ dependencies. A complete list of these dependencies is provided in Table S8.

### Software implementation

NanoTrans is implemented using Perl, Python, R, and Bash scripts. The analysis codes are first encapsulated into functions using Perl, Python or R scripts. The Bash scripts are then used to package these functions along with their required parameters into individual modules. Each module can therefore be executed through one line bash command.

### Hardware requirements

NanoTrans can be deployed on any Linux-based x86-64-bit computing platforms, including cloud service such as Amazon AWS. The minimum hardware requirements for installing NanoTrans are listed below: 2 CPUs, 8 GB RAM, 20 GB of disk space. A minimum of four times the size of hard disk storage in relation to the size of raw fast5 read files is strongly recommended.

### Nanopore DRS datasets reanalyzed in this study

#### Dataset 1

Dataset 1 (BioProject accession: PRJEB39819; Reads accession: ERR4977431) contains admixed reads derived from both the budding yeast and mouse transcriptomes (Tudek et al., 2021). All the reads were used for basecalling and reference-based mapping and only those reads that can be mapped to the budding yeast *Saccharomyces* cerevisiae reference genome (R64-1-1) were used for the downstream analysis. The sequenced yeast RNA was derived from the *S. cerevisiae* strain BY4741 growing on YPDA media under 30 °C and represented a single biological replicate, therefore we named this sample as BY4741.WT.30C.rep1. The sequencing device, sequencing flowcell version, and the sequencing kit used for this dataset are Oxford Nanopore MinION, FLO-MIN106, and SQK-RNA002 respectively.

#### Dataset 2

Dataset 2 (BioProject accession: PRJEB32782; Reads accession: ERR3764352-ERR3764359) contains reads derived from two comparison groups (vir-1 vs. VIR-complemented) of Arabidopsis thaliana samples with four replicates in each group (Parker et al., 2020). The samples from the vir-1 group are mutant defective in the function of the *VIRILIZER* gene, which is an important component of the *Arabidopsis* RNA m^6^A writer machinery. We named these samples as vir1.rep1, vir1.rep2, vir1.rep3, and vir1.rep4 respectively. The samples from the VIR-complemented group all contain a complemented functional version of the *VIRILIZER* gene in combination with a fused Green Fluorescent Proteins (GFP) tag. We named these samples as VIRc.rep1, VIRc.rep2, VIRc.rep3, and VIRc.rep4 respectively. The sequencing device, sequencing flowcell version, and the sequencing kit used for this dataset are Oxford Nanopore MinION, FLO-MIN106, and SQK-RNA001 respectively.

#### Dataset 3

Dataset 3 (BioProject accession: PRJEB39819; Reads accession: ERR5534379-ERR5534382) contains reads derived from two comparison groups (TENT5A^WT^ vs. TENT5A^MUT^) of the cell samples with the human embryonic kidney cell line HEK293 background (Gewartowska et al., 2021). The samples from the TENT5A^MUT^ group express a mutant version of the TENT5A protein with mis-sense mutations in its putative catalytic residues (D139N; D141N). There are two replicates in each comparison group. We named the two samples from the TENT5A^WT^ group as TENT5A.WT.rep1 and TENT5A.WT.rep2, while naming the two samples from the TENT5A^MUT^ group as TENT5A.MUT.rep1 and TENT5A.MUT.rep2. The sequencing device, sequencing flowcell version, and the sequencing kit used for this dataset are Oxford Nanopore MinION, FLO-MIN106, and SQK-RNA002 respectively.

#### Dataset 4

Dataset 4 (BioProject accession: PRJEB44348; Reads accession: ERR9286473 and ERR9286521) contains reads derived from two human cancer cell lines: the lung adenocarcinoma cell line A549 and the leukemia cell line K562 (Chen et al., 2021). We named these two samples as SGNex.A549.rep1 and SGNex.K562.rep1 respectively. The sequencing device, sequencing flowcell version, and the sequencing kit used for this dataset are Oxford Nanopore GridION, FLO-MIN106, and SQK-RNA001 respectively.

#### Dataset 5

Dataset 5 (BioProject accession: PRJCA017047; Reads accession: CRR786043, CRR786046, CRR786061, CRR786077) contains reads derived from two comparison groups (*WRKY7* mutant vs. wildtype) of the common wheat *Triticum aestivum* with two replicates in each group (Chen et al., 2023). The sequencing device, sequencing flowcell version, and the sequencing kit used for this dataset are Oxford Nanopore MinION, FLO-MIN106, and SQK-RNA002 respectively.

### Reference genome and annotation of each dataset

By design, NanoTrans supports the reference genome and annotation file released by Ensembl (https://www.ensembl.org) or its sister sites (e.g., Ensembl Fungi, Ensembl Plants, Ensembl Protists, and Ensembl Metazoa). Therefore, for the analysis of each dataset, we retrieved the corresponding reference genome (in FASTA format) and annotation file (in GTF format) from Ensembl (https://www.ensembl.org). Their downloading Uniform Resource Locators (URLs) are listed in Table S1.

### Benchmarking comparison among NanoTrans, MasterOfPores, and FASTdRNA

Dataset 5 was used to compare the computational performance of NanoTrans (https://github.com/yjx1217/NanoTrans, with the committed number of b3112cc), MasterOfPores (https://github.com/biocorecrg/MOP2, with the committed number of b3d5cfb), and FASTdRNA (https://github.com/Tomcxf/FASTdRNA, with the committed number of 86685fb). For each tool, modules corresponding to the commonly supported functions including Nanopore reads preprocessing (basecalling, reads mapping, and reads counting), RNA modification detection, and poly(A) tail length estimation were executed. This benchmarking test was conducted on a Linux-based server with Intel(R) Xeon(R) Gold 6248R CPU (3.00GHz), 256 GB RAM, and the CentOS operating system (release 7.9.2009). A maximum of 12 threads were used throughout the benchmarking analysis.

## Code and data availability

The NanoTrans software is freely distributed under MIT license at GitHub (https://github.com/yjx1217/NanoTrans).

## CRediT authorship contribution statement

**Ludong Yang**: Data curation, Formal analysis, Investigation, Methodology, Software, Validation, Writing - Original Draft, Writing - Review & Editing. **Xinxin Zhang**: Data curation, Formal analysis, Investigation, Validation, Writing - Review & Editing. **Fan Wang**: Data curation, Formal analysis, Investigation, Validation, Writing - Review & Editing. **Li Zhang**: Resources, Writing - Review & Editing, Funding acquisition. **Jing Li**: Data curation, Formal analysis, Investigation, Resources, Writing - Original Draft, Writing - Review & Editing, Funding acquisition. **Jia-Xing Yue**: Conceptualization, Supervision, Data curation, Formal analysis, Investigation, Methodology, Software, Validation, Writing - Original Draft, Writing - Review & Editing, Funding acquisition.

## Acknowledgements

We thank Dr. Song Gao (Sun Yat-sen University Cancer Center) for inspiring discussion. We thank Dr. Long Wang (Nanjing University) for raw fast5 reads downloading. We also thank the facility support from the Single-Molecule Sequencing Platform at Sun Yat-sen University Cancer Center. This work is supported by National Natural Science Foundation of China (32070592 to JXY, 32000395 to JL, 82272789 to LZ), Guangdong Basic and Applied Basic Research Foundation (2022A1515010717 and 2019A1515110762 to JXY and 2022A1515011873 to JL), Guangdong Pearl River Talents Program (2019QN01Y183 to JXY, 2021QN02Y168 to JL), Guangzhou Municipal Science and Technology Bureau (202102020938 to JL), and Young Talents Program of Sun Yat-sen University Cancer Center (YTP-SYSUCC-0042 to JXY and YTP-SYSUCC-0040 to JL).

