## Supplementary Figure for "NanoTrans: an integrated computational framework for comprehensive transcriptome analysis with Nanopore direct RNA sequencing"

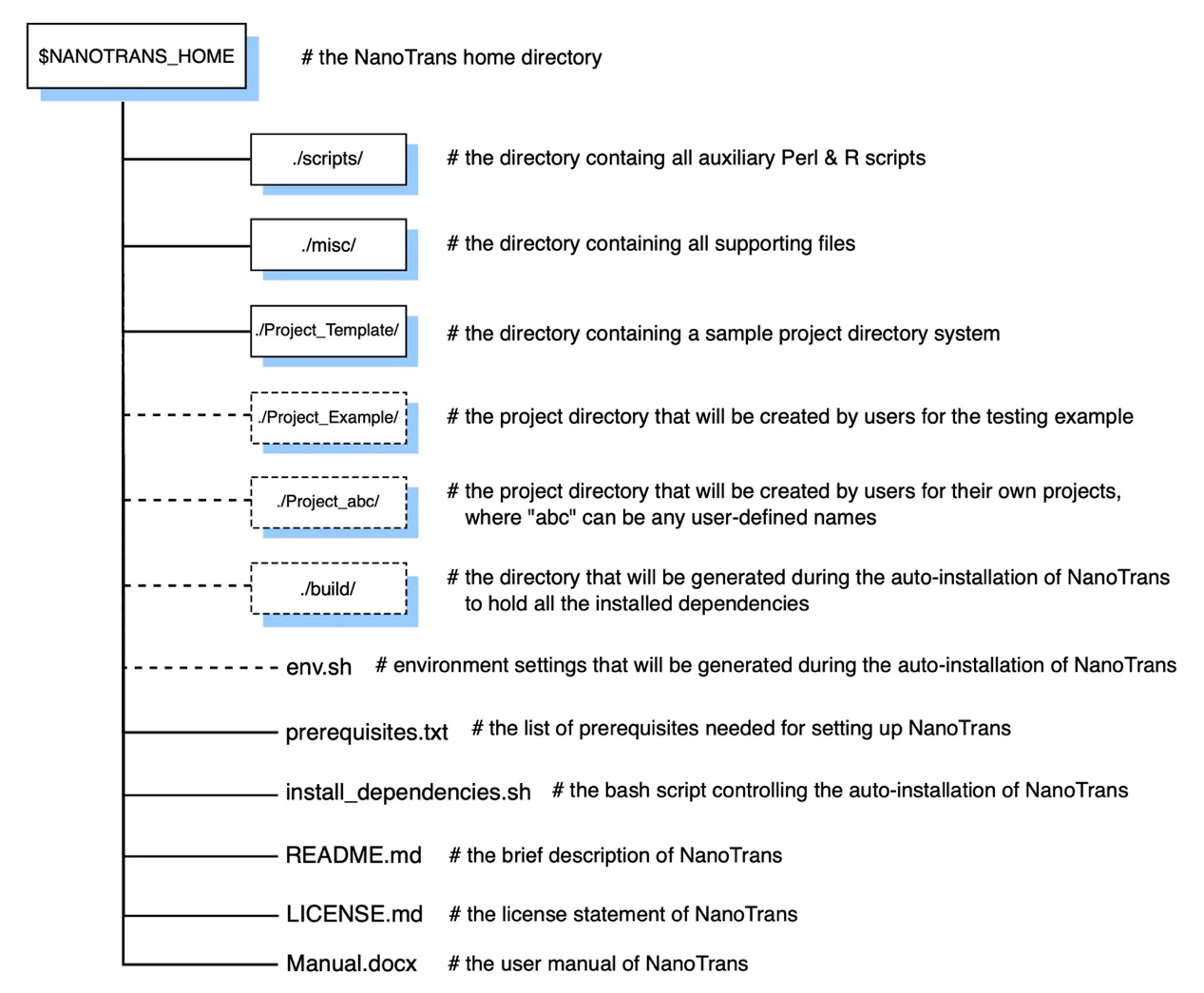

**Figure S1. Overview of the NanoTrans directory system.** All top-level directories (boxes, solid lines) and individual files of NanoTrans are listed and briefly described. Additional directories and files will be generated during the installation and execution of NanoTrans (boxes, dashed lines).

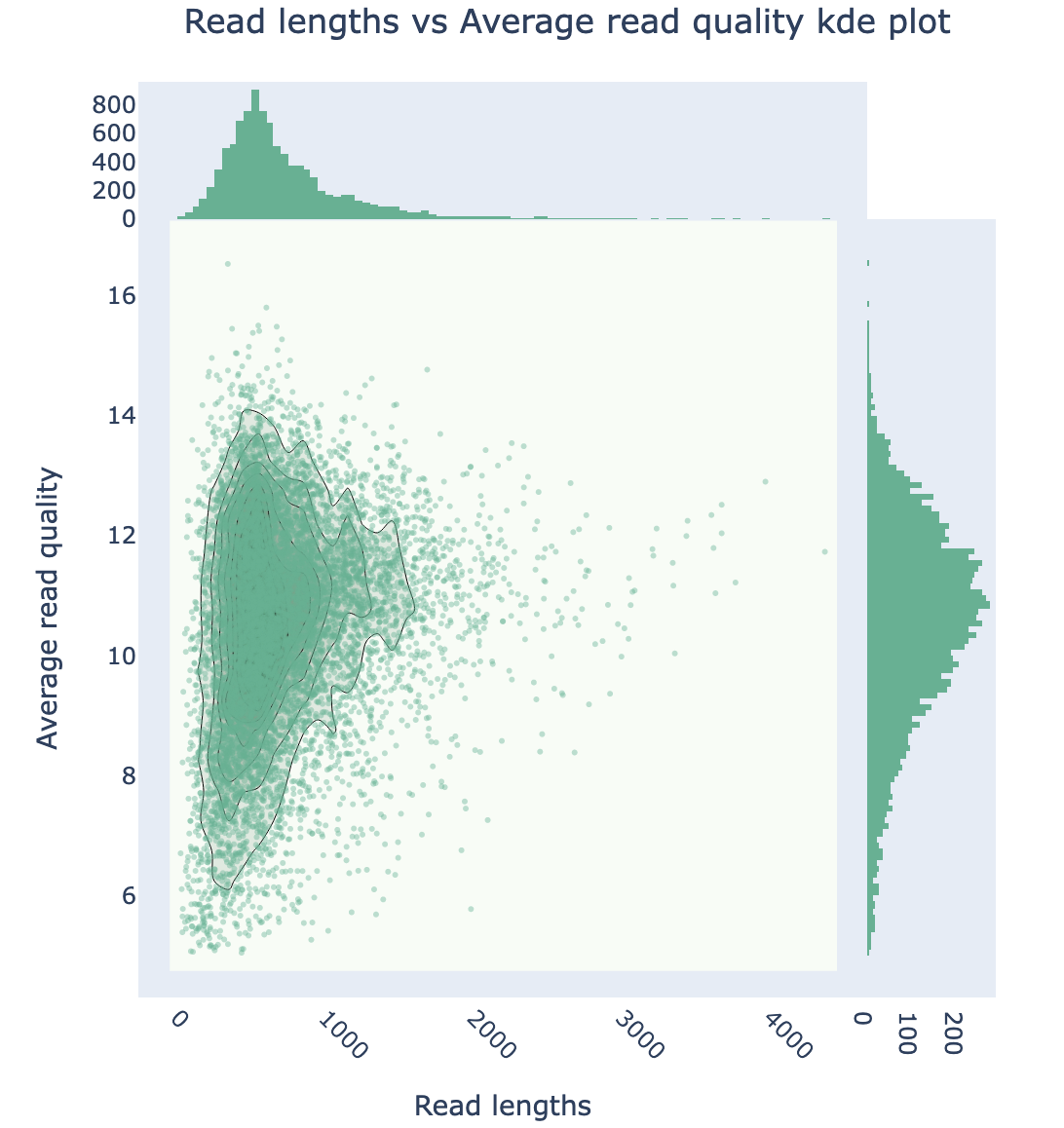

**Figure S2. The read lengths and quality summary plot of basecalled reads from Dataset1.**
